# An integrated data exploration engine of knowledge, expression and genetic association for human transposable elements

**DOI:** 10.64898/2026.10.09.757957

**Authors:** Xiandi Jiang, Yixi Fang, Qiuyu Fang, Shilong Zhang, Guanliang Li, He Xu, Cong Feng, Ming Chen

## Abstract

Transposable elements (TEs) constitute a substantial fraction of the human genome and contribute to genomic and epigenomic regulation. However, TE-related knowledge remains fragmented across the literature and databases, with inconsistent identifiers hindering integration and reuse. Here, we present TE-Dee (Transposable Element Data Exploration Engine), a curated resource that harmonizes literature-derived TE relationships with TE annotations, tissue-specific expression, and GTEx v11 eQTL associations. TE-Dee contains 10,387 entities and 9,583 curated relationships from 2,308 publications, 1,505 leaf-level TE classification records, and 11,334,060 instance-level observations generated by intersecting 640,456 reference TE intervals with significant variant-gene associations across 50 GTEx tissue categories. By linking molecular knowledge to genomic, transcriptomic and genetic evidence, TE-Dee provides reusable data for investigating TE biology, gene-regulatory and disease-related mechanisms, and a structured corpus for training and benchmarking data-driven and artificial intelligence models. TE-Dee is available at https://bis.zju.edu.cn/tedee/.

## Background & Summary

Transposable elements (TEs) are repetitive DNA capable of mobilization or derived from ancestral mobile elements and constitute more than 46% of the human genome^1^. They are commonly classified into retrotransposons, including long interspersed nuclear elements (LINEs), short interspersed nuclear elements (SINEs) and long terminal repeat (LTR) elements, and DNA transposons, together with numerous families and subfamilies that differ in evolutionary age, sequence composition and genomic distribution^2,3^. Although most human TE copies have lost autonomous transposition activity, their sequences remain widely distributed throughout the genome and contribute to genome structure and evolution^4^. TE-derived cis-regulatory elements, such as promoters, enhancers, transcription-factor binding sites, splice sites and polyadenylation signals can influence nearby or distal gene regulation. TE transcription itself varies across cell types, tissues and physiological or pathological states^5,6^. Genetic variants located within TE sequences may further alter these regulatory activities and contribute to inter-individual variation in gene expression^7,8^. Consequently, understanding human TE biology increasingly requires information spanning TE classification, genomic location, transcriptional activity, genetic variation and functional relationships reported in the literature.

A substantial amount of information relevant to human TEs is already available, but it is distributed among resources developed for different purposes. Repeat libraries and genome annotation systems such as Repbase and Dfam provide nomenclature, classification and sequence information for repetitive elements^9,10^. Genome annotations identify individual TE copies at specific chromosomal locations^11^, whereas transcriptomic studies provide measurements of TE activity across tissues and experimental conditions^12,13^. Large-scale human genomics projects, including the Genotype-Tissue Expression (GTEx) project, provide genetic associations between sequence variants and gene expression across multiple tissues^14^. In parallel, experimental and computational studies have reported associations of specific TEs, TE families or TE-derived sequences with genes, biological processes and disease-related phenotypes^15,16^. These data collectively provide complementary views of TE function, but they are typically accessed and analysed separately.

Integration across these resources remains challenging because TE information is represented at several biological and annotation levels. A TE may be described as a broad class, a family or subfamily, a consensus sequence, or an individual genomic insertion. In addition, identifiers and naming conventions are not always consistent between publications, Repbase/Dfam records, RepeatMasker^17^ annotations and downstream genomic datasets. Literature descriptions may use historical names, aliases or different levels of taxonomic resolution, whereas genomic analyses generally operate on coordinate-defined TE instances. Expression datasets and genetic association resources introduce additional tissue, gene and variant identifiers. These differences complicate systematic connection of published TE knowledge with sequence annotations and locus-level molecular evidence, and can limit computational reuse of information assembled from heterogeneous sources.

To address the fragmentation of TE-related data and facilitate their systematic integration and reuse, we developed TE-Dee (Transposable Element Data Exploration Engine), a harmonized resource that integrates literature-curated TE relationships, Repbase/Dfam classifications and sequences, reference-genome TE loci, tissue-specific expression and co-expression profiles, and GTEx v11 expression quantitative trait locus (eQTL) associations (Fig. 1). TE-Dee comprises a knowledge graph of 10,387 entities and 9,583 curated relationships derived from 2,308 publications, together with 1,505 leaf-level Repbase-Dfam classification records. Integration of 640,456 reference TE intervals with significant variant-gene associations across 50 GTEx tissue categories yielded 11,334,060 instance-level observations linking TE-overlapping variants to genes and tissue contexts. Through integrated search, network visualization, genome exploration and expression analysis, TE-Dee connects literature-derived knowledge with genomic, transcriptomic and genetic evidence. Its source-linked, structured datasets facilitate systematic investigation of TE biology and gene regulation and provide reusable resources for computational analysis, model training and benchmarking.

**Fig. 1.**
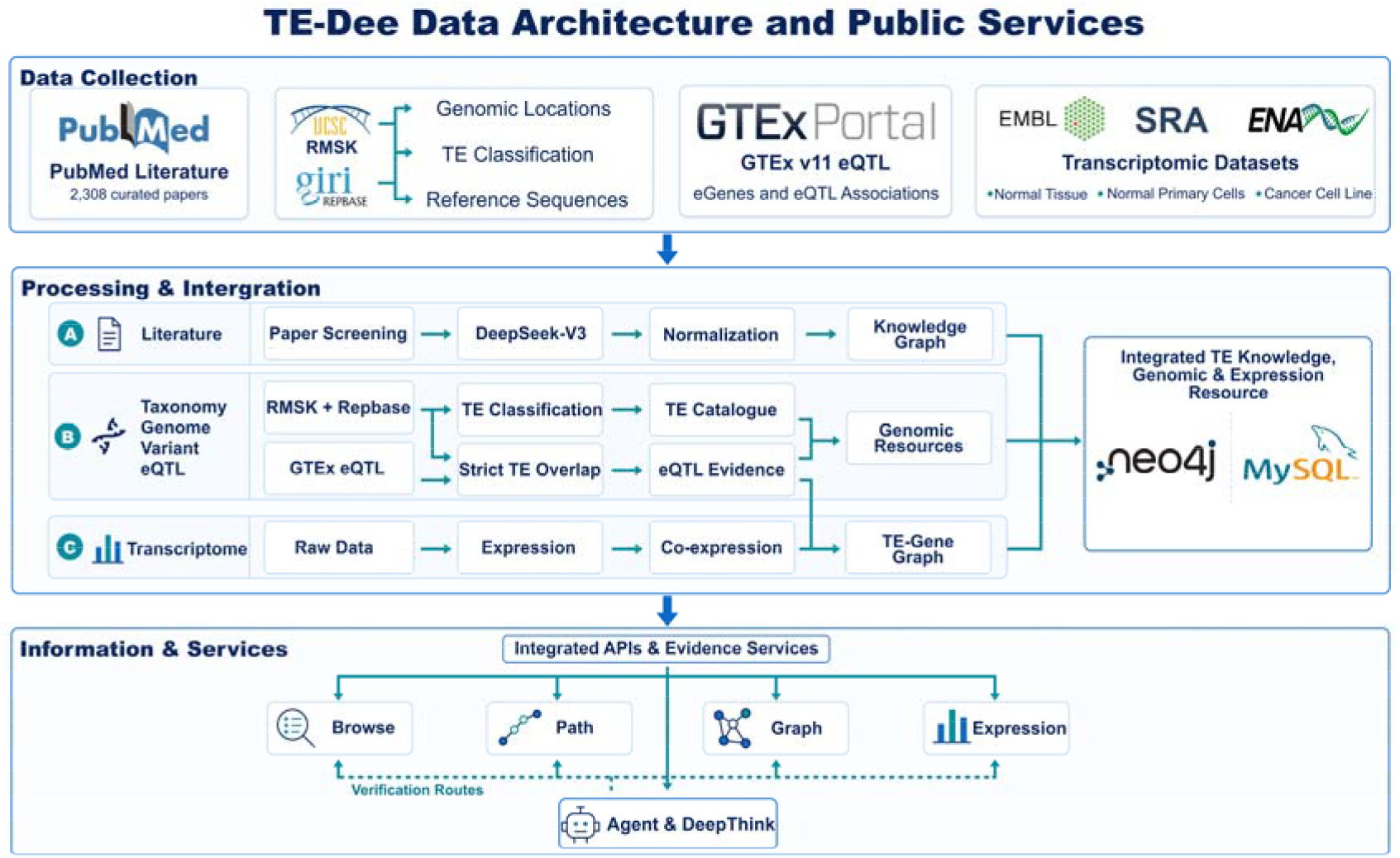
Schematic representation of the TE-Dee data sources, processing and services.

## Methods

### Data sources

TE-Dee was constructed using four categories of data: biomedical literature, TE reference annotations, transcriptomic datasets and genetic association data, supplemented by a standardized gene nomenclature resource.

#### Biomedical literature

Publications served as the primary source for constructing the TE knowledge graph. We retrieved studies related to human TEs from PubMed. The search strategy and subsequent literature processing procedures are described below.

#### TE reference annotations

TE classification, sequence and genomic annotation data were obtained from Repbase/Dfam and UCSC Genome Browser, which provide complementary information at the TE reference and genomic instance levels, respectively. Repbase/Dfam provided TE names, aliases, descriptions, hierarchical classifications and representative sequences. UCSC Genome Browser provided genome-wide annotations of individual TE occurrences, including repeat names, classes, families, chromosomal locations, genomic coordinates and strand orientations.

#### Genetic association data

Single-tissue cis-expression quantitative trait locus (cis-eQTL) data were obtained from the adult GTEx v11 release^18^, which contain tissue-specific variant-gene association records, including variant and gene identifiers, nominal P values and estimated effect sizes (slopes).

#### Transcriptomic datasets

Expression data were collected from three biological contexts: normal tissues, normal primary cells and cancer cell lines. The normal tissue datasets were obtained from ArrayExpress (E-MTAB-1733^19^ and E-MTAB-2836^20^), primary cell data from SRA (SRP013565^21^), and cancer cell line data from BioProject (PRJNA523380), associated with the Cancer Cell Line Encyclopedia (CCLE)^22^. Gene symbols were standardized against the HUGO Gene Nomenclature Committee (HGNC) complete gene set^23^ to ensure consistent gene identification across datasets.

### Literature retrieval and curation

We systematically searched PubMed for publications related to human transposable elements published on or before 13 April 2026. The search strategy combined the MeSH term “DNA Transposable Elements” with the free-text terms “retrotransposon”, “transposon”, “retrotransposons”, “transposons”, “retrotransposition” and “transposition” using the OR operator, together with a human-specific restriction comprising the MeSH term “Humans” and the terms “human” and “Homo sapiens”. The search retrieved 34,788 records. To identify TE-relevant publications, titles and abstracts were screened against a controlled vocabulary compiled from human RepeatMasker repeat names and Repbase identifier and keyword fields. This initial screening excluded 27,118 records, leaving 7,670 publications for structured information extraction.

Entities and relationships were extracted from the retained abstracts using DeepSeek-V3, with a predefined prompt designed to generate structured JSON outputs. The prompt specified the target organism, predefined entity and relationship types, and required PMID identifiers, while restricting extracted descriptions to information explicitly supported by the input abstract. The permitted entity types included TE, disease, disease category, biological function, mutation, gene, RNA, protein, carbohydrate, lipid, peptide, pharmaceutical and toxin. Following extraction, the controlled vocabulary was reapplied to identify records containing human TEs. This second screening excluded 5,362 publications, including 1,803 flagged as potentially lacking a human TE, resulting in a final collection of 2,308 publications. One entity classified as a non-human TE was retained in the resulting knowledge graph.

To harmonize entity nomenclature and reduce redundancy, extracted entity names were grouped within each entity type using the Ratcliff/Obershelp string similarity algorithm with a similarity threshold of 80%. The resulting groups were manually reviewed to establish synonym-to-canonical-name mappings, which were subsequently applied to standardize entity identifiers across publications. The curated entities and their directed relationships were integrated into the knowledge graph, with source PMIDs preserved for each relationship to ensure traceability to the supporting literature.

### TE identity and annotation

TE names, aliases, descriptions, hierarchical classifications and representative sequences were obtained from Repbase-derived records, while genomic repeat annotations and additional classification information were retrieved from the UCSC Genome Browser RepeatMasker (RMSK) track. These complementary sources were harmonized into a unified Repbase-RMSK catalogue to support TE browsing and classification. The catalogue was maintained separately from literature-extracted TE entities, with curated mappings linking original source labels to standardized catalogue names.

Representative or consensus sequences were retained for their corresponding TE entries, whereas individual genomic occurrences were represented by chromosome, start and end coordinates, and strand orientation, allowing a single TE entry to be associated with multiple genomic loci. Disease-related entities were independently organized using an ICD-11-derived classification hierarchy, supplemented with manually curated categories for terms not adequately represented by the selected ICD-11 branches. Disease classifications were assigned on 14 April 2026.

For integration with GTEx eQTL data, TE intervals from the hg38 RMSK annotation were selected through curated mappings between catalogue names and RMSK repeat names. Each selected interval retained its original repeat name, class, family, chromosome, genomic coordinates and strand orientation. Of the 276 approved catalogue names considered for genomic mapping, 226 were successfully matched to hg38 RMSK intervals, while the remaining 50 were retained in the released catalogue without corresponding genomic intervals.

### Taxonomy tree

Two complementary TE taxonomy trees were constructed and stored separately in Neo4j, with the Repbase-RMSK classification rooted at “Transposable Elements - Human” and the broader TE classification rooted at “Mobile genetic element”, which additionally included TE entries absent from RMSK and entries without an explicit taxonomic classification in Repbase. Both trees preserved a hierarchical structure comprising class, subclass, order, superfamily, family and subclade levels. Each taxonomy record retained its original source label, taxonomic rank, hierarchical depth, leaf status and complete classification path. Parent-child relationships were represented independently of the biological relationships extracted from the literature, thereby distinguishing taxonomic organization from reported molecular and functional associations.

Literature-derived TE entities were linked to corresponding taxonomy records through standardized name matching, enabling their classification within the established hierarchies while preserving the distinction between literature-derived knowledge and reference annotations. The Repbase-RMSK taxonomy tree comprised 1,636 records connected by 1,635 parent-child relationships, including 1,505 leaf-level entries. This hierarchy underlies the TE classification browser and classification summaries provided by TE-Dee.

### Expression data processing

Expression datasets were organized into three biological contexts: normal tissues, normal primary cells and cancer cell lines. Raw sequencing reads were subjected to quality control using fastp^24^ and aligned to the GRCh38 human reference genome using STAR^25^. TE read counts were quantified using TEcount from the TEtranscripts suite^26^, incorporating RepeatMasker-based TE annotations. Raw counts were subsequently normalized using the median-of-ratios method implemented in DESeq2^27^.

Expression features were annotated by matching their original identifiers against curated TE reference names and the HGNC complete gene set. Features exactly matching a curated or classified TE name were designated as high-confidence TEs, whereas those exactly matching an approved HGNC symbol were designated as high-confidence genes. Unique matches to previous gene symbols or aliases were assigned lower-confidence gene annotations, while unmatched features remained unassigned. When a feature matched both a TE name and a gene symbol, the TE annotation was prioritized and the ambiguity was recorded. This procedure identified 290 high-confidence TE features and 23,148 high-confidence gene features for subsequent co-expression analysis.

### Co-expression networks construction

Co-expression networks were constructed independently for the three expression settings. Genes were considered eligible if their normalized expression counts exceeded 1 in at least 10% of samples within the corresponding dataset. Expression values were log2-transformed, and eligible genes were ranked by median absolute deviation (MAD), with ties resolved sequentially by variance and feature name. The 2,000 most variable genes were selected alongside the 290 TE features, yielding 2,290 features per dataset.

Pairwise Spearman rank correlations were calculated among the selected features, encompassing TE-TE, TE-gene and gene-gene pairs. Expression values were ranked independently for each feature, with tied values assigned average ranks, and correlations were computed from standardized rank vectors. Features with constant expression, for which correlations were undefined, were excluded from the resulting edge tables. Before correlation analysis, expression matrices were checked for duplicate feature identifiers, missing or non-numeric values, and negative expression counts.

The 2,290 selected features generated 2,620,905 unique unordered pairs per dataset. Significant co-expression relationships were defined by an absolute Spearman correlation coefficient (|r|) ≥ 0.4 and a false discovery rate (FDR)-adjusted P-value ≤ 0.05. Both positive and negative correlations satisfying these criteria were retained, with their correlation coefficients and directions preserved. Co-expression modules were identified from the positive-correlation subgraph using the Louvain community detection algorithm^28^, with correlation coefficients as edge weights, a resolution parameter of 1.8 and a random seed of 42, implemented in NetworkX version 3.4.2. Module assignments and TE-centred subnetworks were stored separately from the complete filtered edge tables.

### eQTL overlap

Single-tissue cis-eQTL data were obtained from the adult GTEx v11 release, based on GENCODE v47 gene annotations. GTEx variant identifiers were parsed to extract chromosome, genomic position, reference allele and alternate allele using GRCh38 coordinates. Variant positions were converted into 0-based, half-open intervals to match the hg38 RepeatMasker annotations. For a variant at 1-based position p, the interval was defined as [p − 1, p − 1 + length(REF)], where REF denotes the reference allele. Variants were considered to overlap a TE instance when they occurred on the same chromosome and their genomic intervals shared at least one base pair. This approach accommodated both single-nucleotide variants and variants with multi-base reference alleles, including those spanning TE boundaries.

Overlapping variants were linked to their corresponding tissue-specific variant-gene associations. Each unique tissue-variant-gene combination was retained as an evidence record, while associations overlapping multiple TE instances were linked to each corresponding interval. Gene annotations, including gene names, biotypes, genomic coordinates and strand orientations, were obtained from the matched eGenes files. Original versioned gene identifiers and association statistics, including nominal P-values, effect sizes, standard errors and allele frequencies, were preserved to maintain traceability to the GTEx source data.

### Data consolidation and integration

Following tissue-specific eQTL processing, the results were consolidated into tables comprising tissues, TE instances, variants, genes, TE-variant overlaps and variant-gene associations, together with tissue-specific and cross-tissue TE-gene summaries. Each TE-variant overlap was stored once and linked to its corresponding associations, preserving variants overlapping multiple TE instances. TE-gene summaries were grouped by TE name, full gene identifier and tissue, or aggregated across tissues. Each summary included the numbers of supporting variants, TE instances and instance-level observations, the minimum nominal P-value and the maximum absolute effect size. Cross-tissue summaries additionally recorded the number of contributing tissues. The resulting tables were exported as gzip-compressed TSV files, accompanied by manifests documenting file structure, record counts and SHA-256 checksums.

To integrate eQTL and co-expression evidence, gene symbols from co-expression datasets were matched to GTEx gene annotations using high-confidence, exact and unambiguous identifiers. TE-gene pairs supported by both a retained co-expression relationship and TE-overlap eQTL evidence were classified as “Both”, whereas pairs supported by only one evidence type were labelled “Co-expression” or “eQTL”. Expression settings and GTEx tissue contexts were retained separately to preserve their respective biological contexts.

### Data services

TE-Dee provides a web interface for human TE classification, reference sequences, genomic annotations, literature-derived relationships, expression profiles and eQTL evidence. Browse presents available sequences and genomic records for individual TEs, with genomic visualization through JBrowse 2^29^. Graph and Path display relationships among TEs, genes, diseases and biological functions alongside supporting publications. Expression summarizes TE abundance across normal tissues and normal and cancer cell lines.

The TE-Gene Graph combines context-specific co-expression with GTEx eQTL evidence from individual tissues or across tissues. TE-gene pairs are labelled as supported by co-expression, eQTL associations or both. The Integrated Graph combines these networks with literature-derived relationships and retains their evidence types and provenance. Enrichr^30^ tests gene sets from the network for enrichment in GO biological processes and KEGG pathways.

Agent and DeepThink answer natural-language questions using the available evidence and retain conversational context for follow-up questions. The Download section provides datasets for further analysis and reuse.

### Data Records

The records are available from the download page (https://bis.zju.edu.cn/tedee/download.php) and are organized in five layers: literature, TE reference, expression, co-expression and eQTL. Each layer is stored as its own set of files, and the TE catalogue name connects the layers where a match has been established.

TE-Dee contains 1,505 TE catalogue entries, 10,387 entities and 9,583 relationships (Fig. 2a-c). ERV1 (382 entries) and ERV3 (HERV-R; 244 entries) were the largest TE superfamilies. Function was the largest entity category, with 3,683 entities. Of the relationships, 1,635 represented taxonomic classification, while the remainder covered biological associations. The genomic tracks mapped TE copies, co-expressed genes and TE-overlapping eQTL variants across GRCh38. Their distributions varied across genomic regions, and the co-expression profiles differed among normal tissues, primary cells and cancer cell lines (Fig. 2d).

**Fig. 2.**
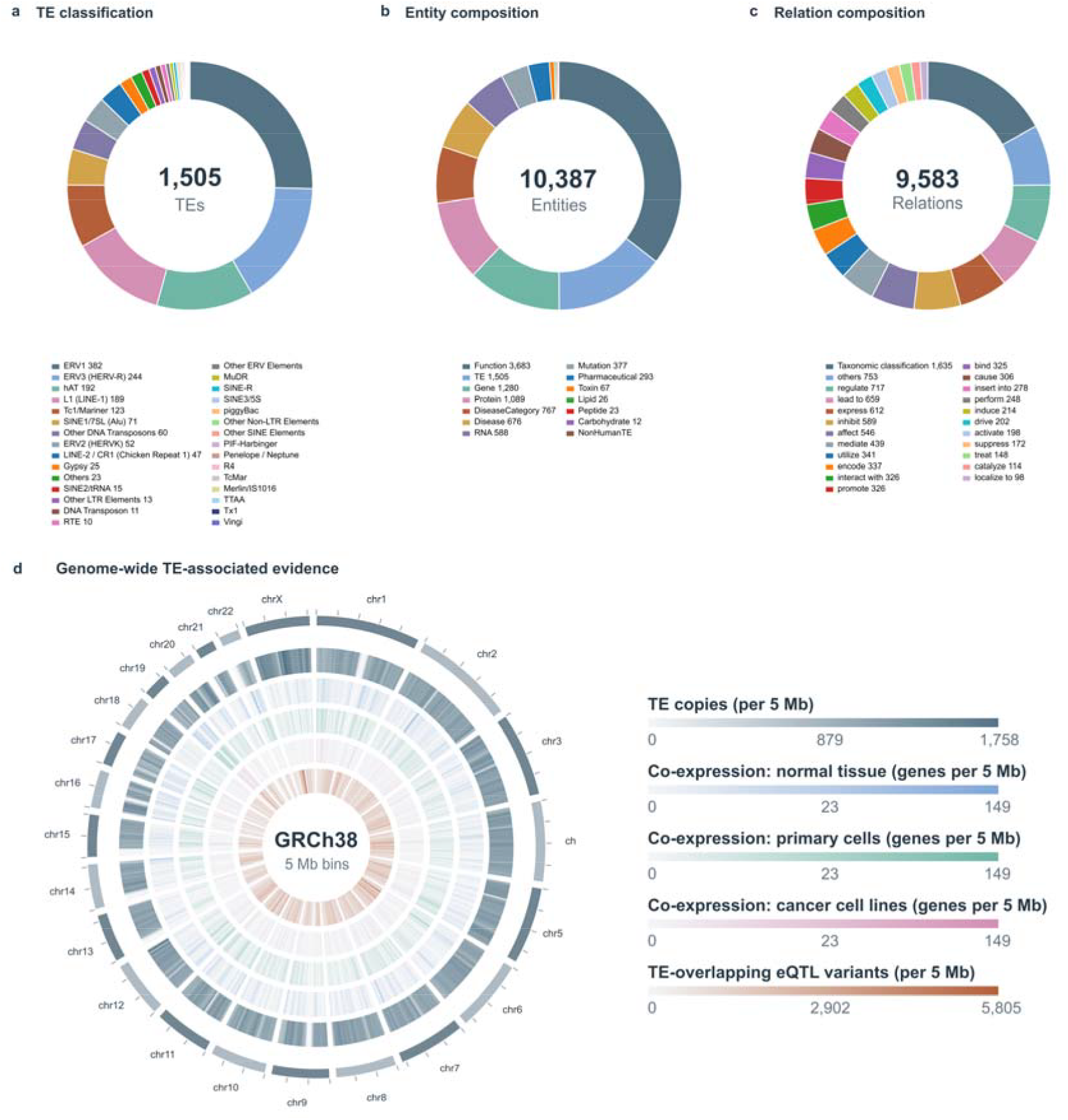
Composition of the TE-Dee records and the genome-wide distribution of TE-associated evidence. (a-c) The Repbase + RMSK classification catalogue, the entity types and the relation types. (d) Density of TE copies, genes co-expressed with TEs, and TE-overlapping eQTL variants in 5-Mb bins across chromosomes 1-22 and X. Colors represent the density, as reported in the legend.

### Literature records

Publications, typed entities and directed relationships are stored as separate tables. A relationship row gives the source entity, the predicate, the target entity and the PMID or PMIDs of the supporting articles. Entity rows carry the entity type, the curated name and the recorded aliases. The tables use the identifiers of the graph database, and the release contains the extracted records.

### Catalogue and genomic records

The catalogue contains 276 entries, each with a name, aliases, classification path and source label. The two taxonomy trees are supplied as node and parent-child tables; each node retains its source label, rank, depth, leaf status and complete path. Genomic records contain one row per hg38 RepeatMasker occurrence, giving chromosome, start, end, strand, source repeat name, class, family and any mapped catalogue name. The reference annotation file provides representative or consensus sequences. These genomic occurrences underlie the TE-copy density track in Fig. 2d.

### Expression records

Each of the three expression settings is provided as a tab-separated matrix with features in rows and samples in columns, together with a sample metadata table that assigns every column to a biological setting. Additional tables contain the context summaries and the identity and annotation confidence assigned to each feature. Fig. 3 shows the relative expression of classified TE superfamilies across the 89 contexts.

**Fig. 3.**
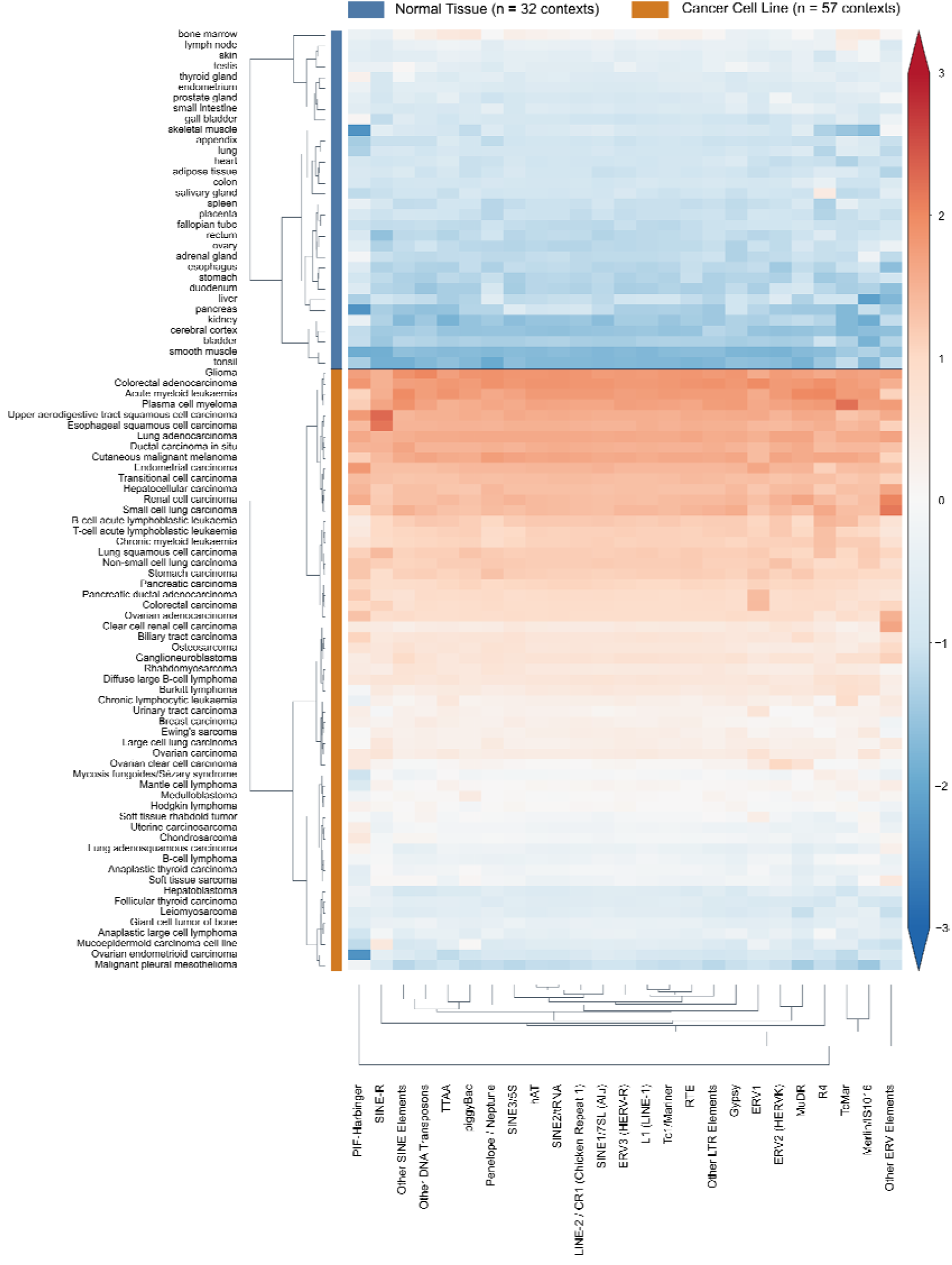
Relative expression of TE superfamilies across normal-tissue and cancer-cell-line contexts. Expression values of taxonomy-matched TE leaves were summed within each superfamily and context, transformed as log2(sum + 1), and converted to a Z-score across the 89 contexts. Colors represent the Z-score, as reported in the legend.

Normal-tissue contexts generally showed negative Z-scores, while positive scores were common among cancer-cell-line contexts. Most cancer-cell-line contexts, including glioma and colorectal adenocarcinoma, showed positive Z-scores across all 25 displayed superfamilies. Malignant pleural mesothelioma and ovarian endometrioid carcinoma showed negative scores across the same superfamilies, illustrating variation within the cancer-cell-line group.

### Co-expression records

The co-expression files contain the selected features, the complete filtered edge table, the module assignments of the positive-correlation network and the TE-centred subgraphs used by the web view. An edge records the two feature identifiers and their types, the expression setting, the correlation, the adjusted p-value and the number of samples. The selection table records the decision and the reason for every candidate feature.

### eQTL records

Each GTEx tissue directory contains te_variant_gene_overlaps.parquet, te_gene_summary.parquet and a processing record. Eight consolidated tables are also available as gzip-compressed TSV files. Table 1 gives the record unit and identifying fields for each table. An instance-variant overlap is unique by instance and variant keys, while an association is unique by tissue, variant and full gene identifier. The identifier without a version suffix is provided only for the checked match to co-expression gene symbols. Joining the overlap and association tables by variant key recovers evidence for an individual TE instance. Each file’s processing record lists its name, field order, byte size and checksum.

**Table 1.**
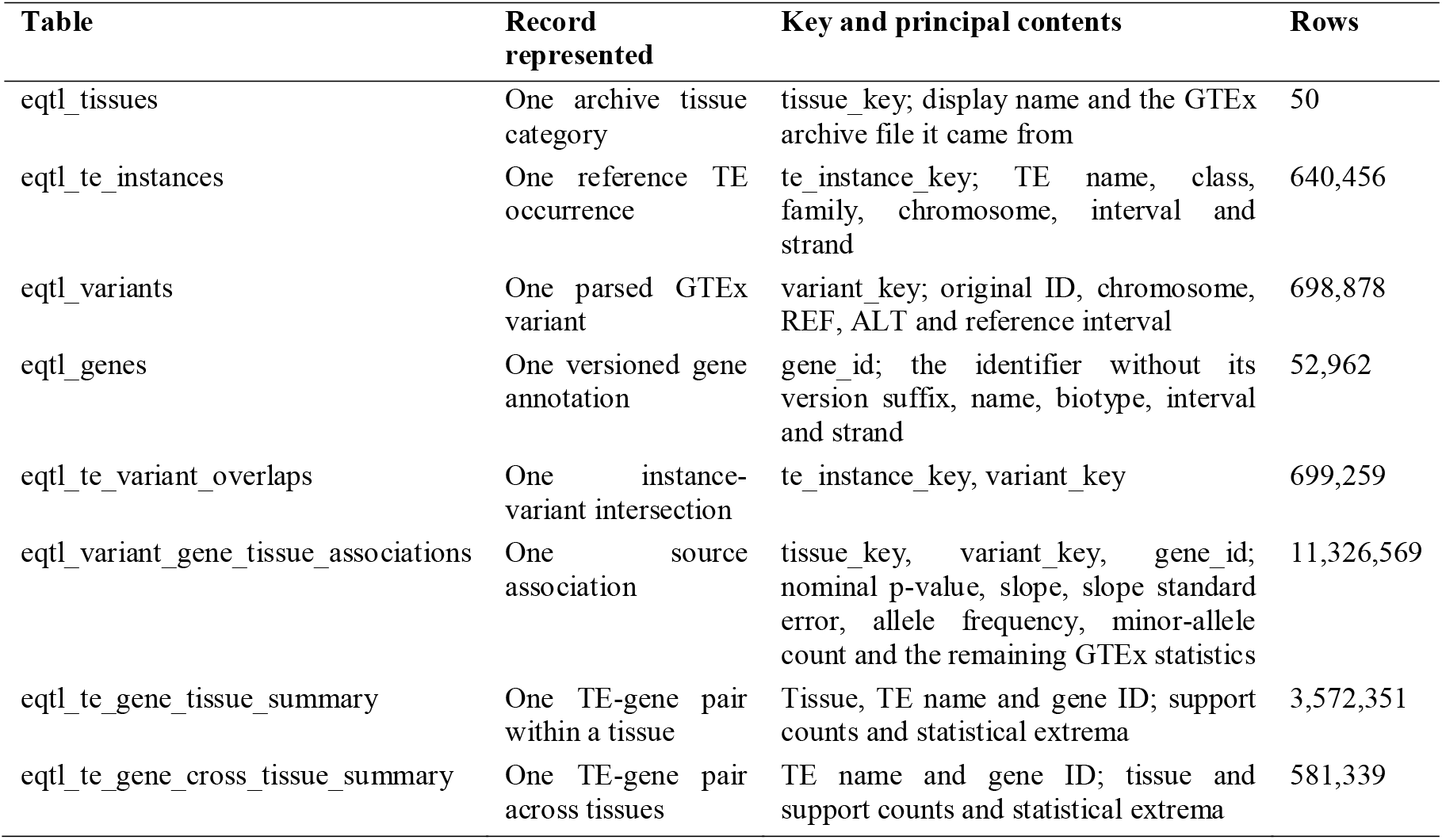
Normalized eQTL records and their identifying fields.

The eight tables contain 17,571,864 rows in total. The GTEx v11 association files are available from the GTEx download collection ^18^.

## Technical Validation

### Literature screening

We sampled three sets of records to assess screening decisions: those removed in the first screening, those removed in the second screening and those flagged during graph integration. Fifty records were drawn at random from each set. Manual review judged 50, 47 and 48 decisions correct, respectively.

### Identifier and record consistency

We checked feature annotations and the gene identifiers linking co-expression results to GTEx eQTL tables. TE-gene co-expression edges contained 3,281 distinct gene symbols. Among them, 3,243 had a high-confidence co-expression annotation, an exact GTEx gene-name match and a unique base gene identifier. The remaining 38 had no exact gene-name match in the GTEx annotation.

Across the three contexts, the audit counted 170,599 TE-gene co-expression records representing 169,474 distinct TE-gene pairs. An eligible unique high-confidence mapping was available for 167,997 records; 2,602 lacked one. Among eligible records, 7,763 had cross-tissue eQTL evidence for the same TE-gene pair and 160,234 did not. Before tissue or display filtering, 7,715 distinct pairs carried both evidence types.

The audit matches symbols by exact case and requires a single identifier after version suffixes are removed. These are the rules of the audit; the interface resolves the same symbols case-insensitively against a list of high-confidence gene names.

We also compared the three expression matrices. They contained the same feature set and no duplicated feature label, and every TE or gene feature selected for co-expression was present in the corresponding matrix. Sample columns were matched to their metadata before the context summaries were calculated; of the 205 normal-tissue columns, 200 could be linked and were used for the summaries. For the literature layer, each released relationship carries a predicate and the PMID or PMIDs of the articles that support it.

### Overlap and consolidation

We tested the interval procedure using synthetic variant and TE intervals with known outcomes. Ten cases covered single-nucleotide and multibase reference alleles, chromosome labels, boundary contact without a shared base and variants overlapping multiple TE instances. Indexed interval-search results agreed with direct comparisons of the same intervals. The cases also tested tissue-file pairing, annotation joins, duplicate removal, conflict detection and repeatable export.

We reconciled counts from tissue processing with the consolidated tables. Tissue outputs contained 104,901,807 source association rows and 11,334,060 instance-level observations. The eight consolidated tables contained 17,571,864 rows, including 11,326,569 in the association table. Direction classes in the summary tables derive from the signs of contributing slopes. The catalogue mapping accounts for all 276 entries; unmatched names remained unmatched.

### Co-expression construction

Seven checks covered feature selection, pair construction, multiple-testing adjustment, edge filtering and community detection, and all passed. The processing record of each network states the transformation, the gene-selection settings, the number of tested pairs, the handling of positive edges, the software version, the resolution and the random seed, so each network can be rebuilt from the released node and edge tables together with the selection table.

### External comparison and limitations

Two checks compared our records with external references. Against the GTEx gene annotation, 3,243 of 3,281 co-expression gene symbols (98.8%) resolved to a unique identifier after version suffixes were removed. The 38 unmatched symbols fell into four groups: nine Y-chromosome genes (DDX3Y, EIF1AY, KDM5D, NLGN4Y, PRKY, RPS4Y1, TMSB4Y, TTTY10, UTY); twelve antisense or long intergenic non-coding RNA symbols, including ANO1-AS1 and LINC01422; six immunoglobulin or T-cell receptor segments (IGHJ3P, IGHV1-24, IGHV1-69-2, IGLC1, IGLJ1, TRBJ2-2P); and eleven other symbols without an exact match in the GTEx v11 annotation.

The second comparison is the coverage of the catalogue. Of the 276 catalogue names, 226 (82%) were matched to hg38 RepeatMasker intervals and 50 were not, and only matched intervals entered the overlap analysis. The eQTL layer therefore covers the part of the catalogue that the mapping placed on the reference genome.

### Case study

HSMAR2 was used to illustrate how TE-associated gene networks can be examined through functional enrichment. Fig. 4 shows the ten lowest adjusted P values for the 73 normal-tissue genes connected to HSMAR2. Relationship types and source records document the TE-gene connections underlying this analysis. The GO biological process and KEGG pathway results summarize the functional annotations represented in this gene set and provide candidates for further investigation. The same records are available through a natural-language interface. The service follows the FAIR principles for findable, accessible, interoperable and reusable data ^31^.

**Fig. 4.**
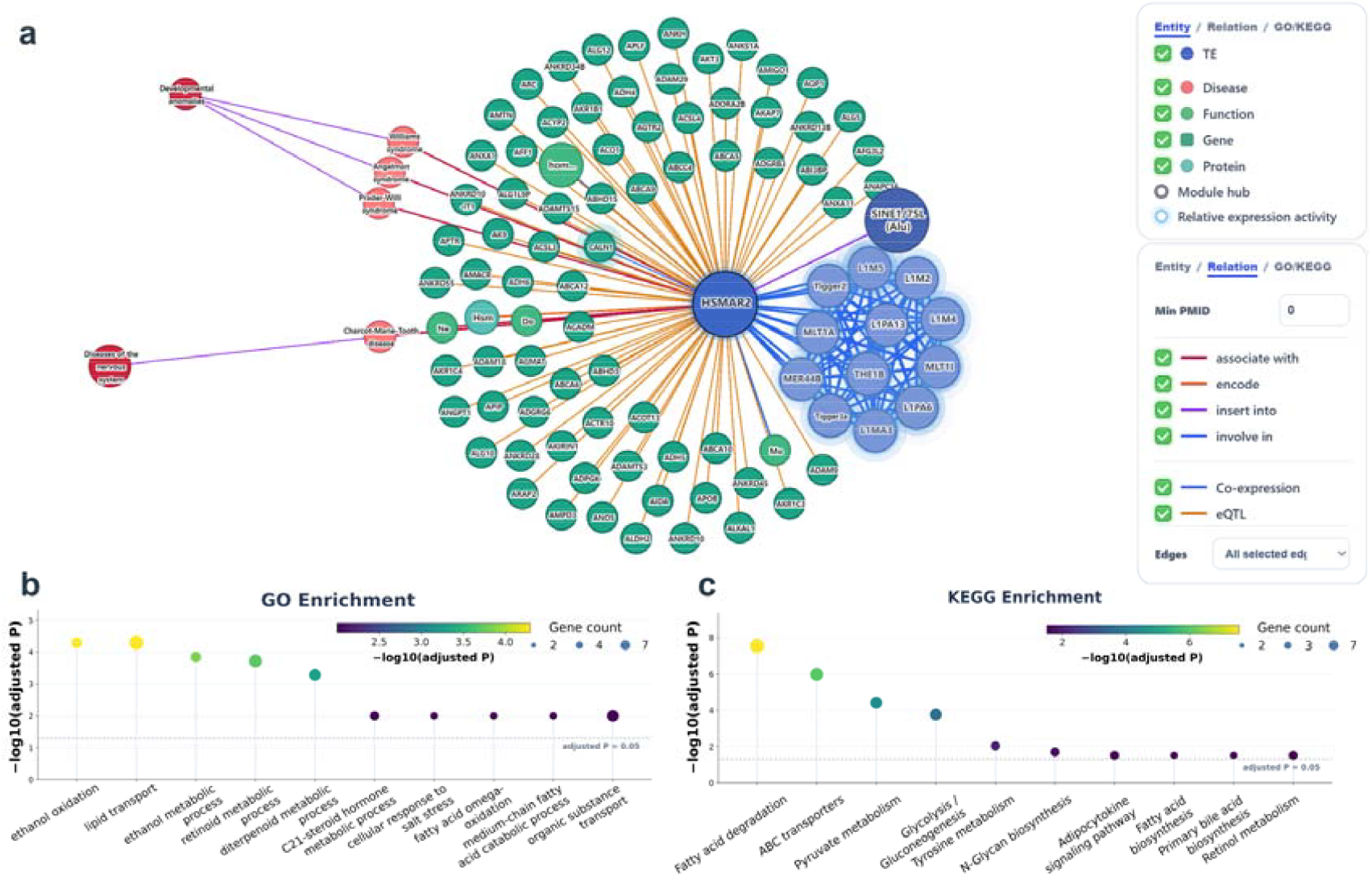
Integrated Graph view of TE-Dee, centred on the TE HSMAR2. (a) The Integrated Graph combines the literature-derived knowledge graph and the TE-Gene Graph in one network; edges are labelled Co-expression, eQTL, or Both. (b,c) Gene Ontology (b) and KEGG (c) terms enriched among the normal-tissue genes connected to HSMAR2; the dashed line marks an adjusted P value of 0.05.

### Usage Notes

The records can be reached by TE name, by biological entity, by taxonomy path, or by publication. From a TE record the interface opens the reference annotation, the classification path, the representative sequence, the genomic occurrences, the expression summaries and the available literature.

Four keys connect the released tables: variant_key identifies a parsed GTEx variant, tissue_key an archive tissue category, gene_id a full versioned gene identifier, and te_instance_key an annotated genomic occurrence. Analyses should specify whether they count associations or instance-level overlaps. One association can intersect multiple TE instances and therefore appears more than once in the instance-expanded table.

The downloadable files contain the complete filtered co-expression networks and the complete tissue tables, whereas the web views show smaller subsets for inspection. Analyses that depend on the full network should use the downloaded tables.

The eQTL layer includes only catalogue names mapped to reference-genome intervals. The 50 names without an hg38 RepeatMasker interval therefore have no overlap records. Expression context summaries include the 200 normal-tissue columns linked to metadata; the five unlinked columns remain in the downloadable matrix. Both limits are documented in the release.

## Data Availability

TE-Dee records are deposited in figshare under DOI 10.6084/m9.figshare.33945424 and are also available from https://bis.zju.edu.cn/tedee/. The website provides the download page for the files described in Data Records. Source expression datasets have accessions E-MTAB-1733, E-MTAB-2836, SRP013565 and PRJNA523380 ^19,20,22^. GTEx v11 association files are available from the GTEx download collection ^18^. No individual-level GTEx data are redistributed.

## Code Availability

The code used in this study is available at https://github.com/DeeJ6126/TE-Dee.git. The repository contains scripts for literature name mapping, feature annotation, co-expression construction, TE-variant overlap, tissue consolidation and website queries. Each data product’s processing record identifies the script and parameters used to produce it.

## Author Contributions

Conceptualization, M.C. and C.F.; methodology, X.J. and Y.F.; software, X.J., Y.F. and S.Z.; validation, Y.F.; formal analysis, X.J.; investigation, X.J. and Y.F.; resources, C.F., G.L. and H.X.; data curation, X.J. and Y.F.; writing—original draft preparation, X.J.; writing—review and editing, C.F. and Q.F.; visualization, X.J.; supervision, M.C. and C.F.; project administration, C.F., X.J. and Y.F. contributed equally. All authors read and approved the final manuscript.

